# Development and Characterization of Recombinant ADP-Ribose Binding Reagents that Allow Simultaneous Detection of Mono and Poly ADP-Ribose

**DOI:** 10.1101/2024.05.16.594588

**Authors:** Shu-Ping Chiu, Cristel V. Camacho, W. Lee Kraus

**Affiliations:** Laboratory of Signaling and Gene Regulation, Cecil H. and Ida Green Center for Reproductive Biology Sciences, University of Texas Southwestern Medical Center, Dallas, TX 75390, USA; Section of Laboratory Research, Department of Obstetrics and Gynecology, University of Texas Southwestern Medical Center, Dallas, TX 75390, USA

**Keywords:** ADP-ribose, ADP-(ribosyl)ation, recombinant fusion proteins, antibody-like reagents

## Abstract

ADP-ribosylation (ADPRylation) is a post-translational modification (PTM) of proteins mediated by the activity of a variety of ADP-ribosyltransferase (ART) enzymes, such as the Poly (ADP-ribose) Polymerase (PARP) family of proteins. This PTM is diverse in both form and biological functions, which makes it a highly interesting modification, but difficult to study due to limitations in reagents available to detect the diversity of ADP-ribosylation. Recently we developed a set of recombinant antibody-like ADP-ribose binding proteins, using naturally occurring ADPR binding domains (ARBDs) that include macrodomains and WWE domains, that have been functionalized by fusion to the constant “Fc” region of rabbit immunoglobulin. Herein, we present an expansion of this biological toolkit, where we have replaced the rabbit Fc sequence with two other species, the Fc for mouse and goat immunogloblulin. Characterization of the new reagents indicates that they can be detected in a species-dependent manner, recognize specific ADP-ribose moieties, and excitingly, can be used in various antibody-based assays by co-staining. The expansion of this tool will allow for more multiplexed assessments of the complexity of ADPRylation biology in many biological systems.

## Introduction

In recent years, there has been a substantial increase in our understanding of the biochemistry, molecular biology, normal physiology and pathology of ADP-ribosylation (ADPRylation) of proteins *1-3*. It is clear that this complex post-translational modification (PTM) plays important roles in many biological processes, including DNA repair, transcription, immune regulation and condensate formation (among many others). This PTM is mediated by a variety of ADP-ribosyltransferase (ART) enzymes, including the Poly (ADP-ribose) Polymerase (PARP) family of proteins (with exception of non-catalytic members). These enzymes mediate the transfer of ADP-ribose from nicotinamide adenine dinucleotide (NAD^+^) to a surprisingly diverse set of substrate proteins, on a wide range of amino acids. The susbtrates can be modified by ADP-ribose chains of varying lengths and chemical structures, i.e. by either a single ADP-ribose unit (monoADPRylation or MAR) or polymers of ADP-ribose units (polyADPRylation or PAR), which serve to alter biochemical activities of the substrate protein or drive protein-protein interactions through new interaction surfaces *1-5*.

ADPRylation “reader” proteins also exist in nature, with modules that can specifically recognize and bind to various forms of ADP-ribose moieties. Some of the most well-characterized ADP-ribose binding domains (ARBDs) include WWE domains, which recognize PAR (and oligo chains) and macrodomains, which recognize MAR or terminal ADP-ribose moieties found in PAR (and oligo chains) *[1, 3, 4, 6]*. Recently we developed a set of recombinant antibody-like ADP-ribose binding proteins, using naturally occurring macrodomains and WWE domains, that were functionalized by fusion to the constant “Fc” region of rabbit immunoglobulin. Our ARBD-Fc fusion proteins represent one example of tools developed for the detection of ADP-ribose moieties. Other examples include the development of monoclonal *7* and polyclonal *8-15* antibodies to detect MAR and PAR, SpyTag-based modular antibodies to detect MAR *16, 17*, and the use of the hydrolases ARH3 and TARG1 as tools to investigate ADPRylation specifically on serine and glutamate residues, respectively *18, 19*. These reagents have all proven useful research tools in the field for the molecular recognition of various forms of ADP-ribose *4*, but significant limitations still exist. Given the complex nature of this modification, there is a need for the continued development of tools and refinement of existing ones, which will undoubtedly enhance our understanding of the intricate biological functions of ADPRylation.

Herein, we describe the generation and characterization of an expanded set of antibody-like ADP-ribose binding proteins previously reported *4*, in which these natural and specific ARBDs have been functionalized with the Fc region of mouse and goat immunoglobulin to create a more useful array of ADP-ribose detection reagents. Characterization of the new reagents indicates that they can be detected in a species-dependent manner, recognize specific ADP-ribose moieties, and excitingly, can be used in various antibody-based assays by co-staining. The expansion of this tool will allow for more multiplexed assessments of the complexity of ADPRylation biology in many biological systems.

## Results

### Expression and purification of new recombinant mouse and goat ARBD-Fc fusion proteins

We previously described the generation of Fc fusion proteins using the macrodomain from *H. sapiens* PARP14 (Macro2 and Macro 3; M2/3), *A. fulgidus* AF1521, and WWE domain from *H. sapiens* RNF146, fused the Fc region of rabbit IgG *4*. We constructed six new vectors by replacing the rabbit-Fc with cDNA from mouse and goat Fc (Figure 1A, Figure S1 and S2). We expressed the ARBD-Fc fusion proteins *in E. coli* and purified them using Ni-NTA affinity chromatography as previously described (Table S1-S2). Purified proteins were analyzed by denaturing SDS-PAGE and stained with Coomassie (Figure 1B; *left panel*). All fusion proteins migrated at the expected molecular weights (Figure 1B; *right panel*). We observed that the new mouse and goat preparations of WWE had high purity, whereas some contaminating bands were present in the M2/3 (PARP14) and AF1521 preps, with a major contaminating band at ∼30 kDa, which did not seem to affect detection (see below).

**Figure 1.**
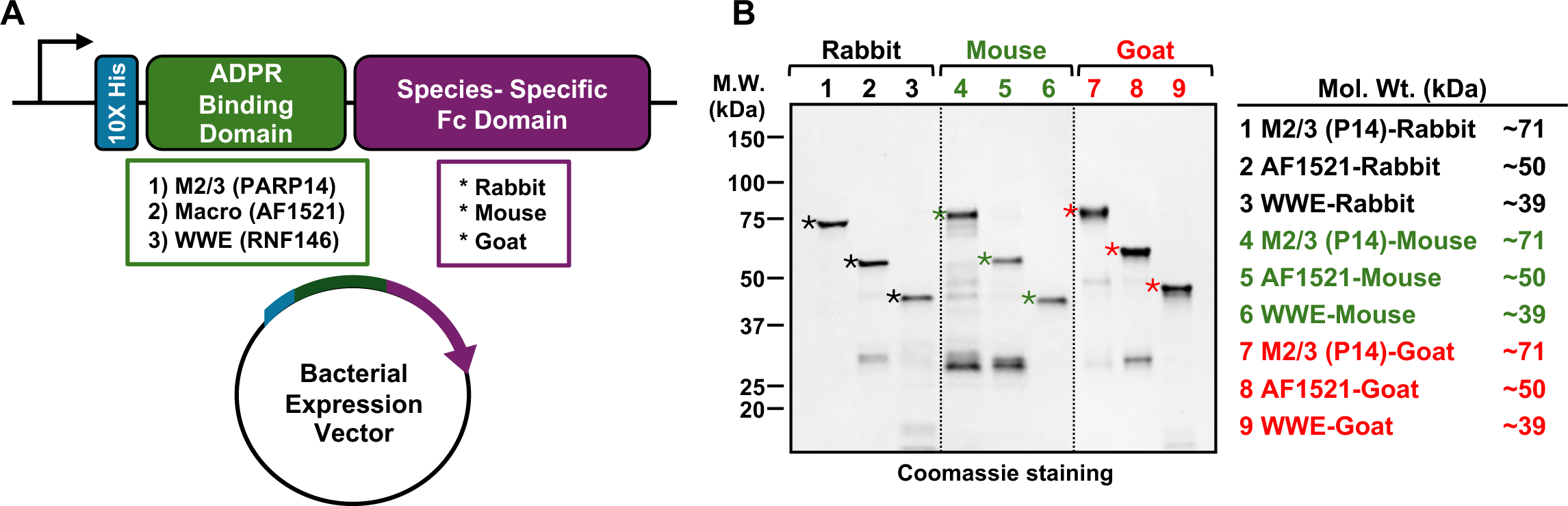
Design, expression, and purification of new ARBD-Fc detection reagents. **(A)** Schematic diagram of the plasmid constructs used to express the ADP-ribose binding domain-Fc (ARBD-Fc) fusion proteins in bacteria. The constructs contain DNA segments encoding (1) 10X His tag (blue), (2) an ADP-ribose binding domain [Macro 2/3 (PARP14), Macro (AF1521), or WWE (RNF146); green], and (3) the rabbit, mouse, or goat IgG constant fragment (Fc; purple). **(B)** Expression and purification of ADP-ribose binding domain-Fc (ARBD-Fc) fusion proteins. ARBD-Fc fusion proteins were expressed in *E. coli* and purified using Ni-NTA agarose affinity resin. The purified proteins were separated by SDS-PAGE and stained with Coomassie brilliant blue (*left panel*). The asterisks indicate protein bands with the expected molecular weights of the ARBD-Fc fusion proteins. Molecular weight markers in kilodaltons (kDa) are indicated. The table is a list of 9 ARBD-Fc fusion proteins and their expected molecular weights (*right panel*).

### ARBD-Fc fusion proteins can be detected in a species-specific manner

To test the ability of the new ARBD-Fc fusion proteins to be detected in a species-specific manner, we carried out dot blots using equal amounts of all nine purified fusion proteins and IgG from rabbit, mouse, and goat as controls (Figure 2A; *left panel*), or SDS-PAGE analysis (Figure 2B). Using species-specific HRP-conjugated secondary antibodies for detection of proteins, we show that each species of ARBD-Fc fusion proteins is detected exclusively by a single corresponding secondary antibody, indicating highly specific detection (Figure 2A-B).

**Figure 2.**
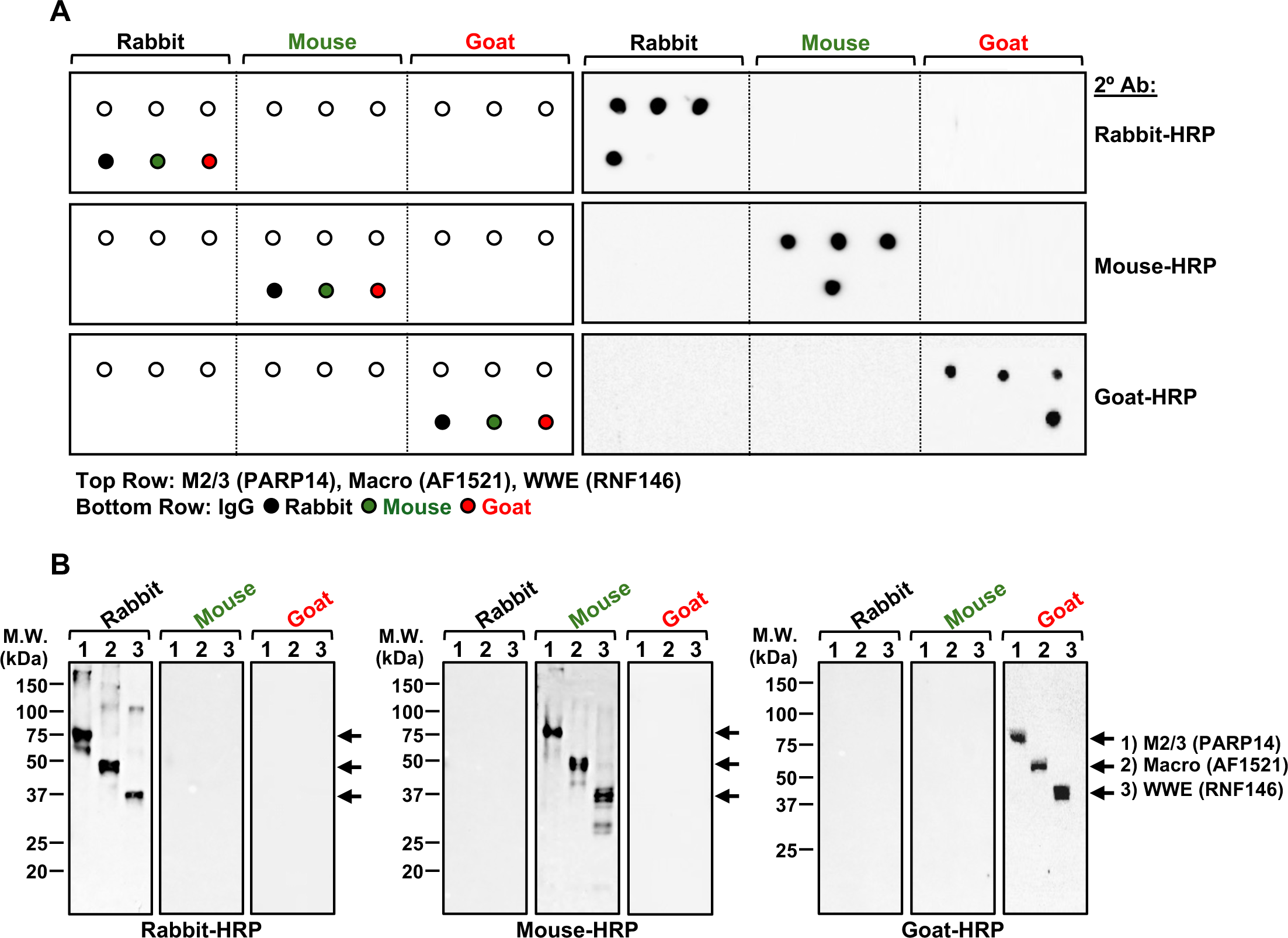
Fc species-specificity of new ARBD-detection reagents. **(A)** Schematic of dot blot (*left panel*) and immunoblot analyses (*right panel*) of species specificity of rabbit, mouse, and goat Fc using ARBD-Fc fusion proteins. Nine purified ARBD-Fc fusion proteins, rabbit, mouse, and goat IgG protein were applied to nitrocellulose membrane for dot blotting (*left panel*). Each spot contained approximately 500 ng of protein and was blotted using anti-rabbit, anti-mouse, and anti-goat HRP-conjugated antibody, as indicated (*right panel*). **(B)** Immunoblot analyses of nine purified ARBD-Fc fusion proteins separated by SDS-PAGE, transferred to nitrocellulose membrane, and subjected to immunoblotting using the anti-rabbit, anti-mouse, and anti-goat HRP-conjugated antibody, as indicated. Molecular weight markers in kilodaltons (kDa) are indicated.

### ARBD-Fc fusion proteins can recognize specific forms of ADPR

To test the binding of the new ARBD-Fc fusion proteins and ability to recognize exact forms of ADPR, we performed in vitro ADPRylation reactions to generate mono ADP-ribose (MAR) or poly ADP-ribose (PAR) chains. As previously described, purified recombinant PARP3 or PARP1 were used in biochemical reactions with NAD^+^ (ADPR donor) and sonicated salmon sperm DNA (activator) to generate automodified PARP3 (MAR) and PARP1 (PAR). Reactions lacking NAD^+^ were used as a control. As expected, in each species version, M2/3 (PARP14) was able to detect only MAR, WWE was able to detect PAR, while AF1521 was able to detect both forms of ADPR (Figure 3A). These results confirm that each domain is able to detect a specific form of ADPR and exchanging the Fc region did not affect this detection.

**Figure 3.**
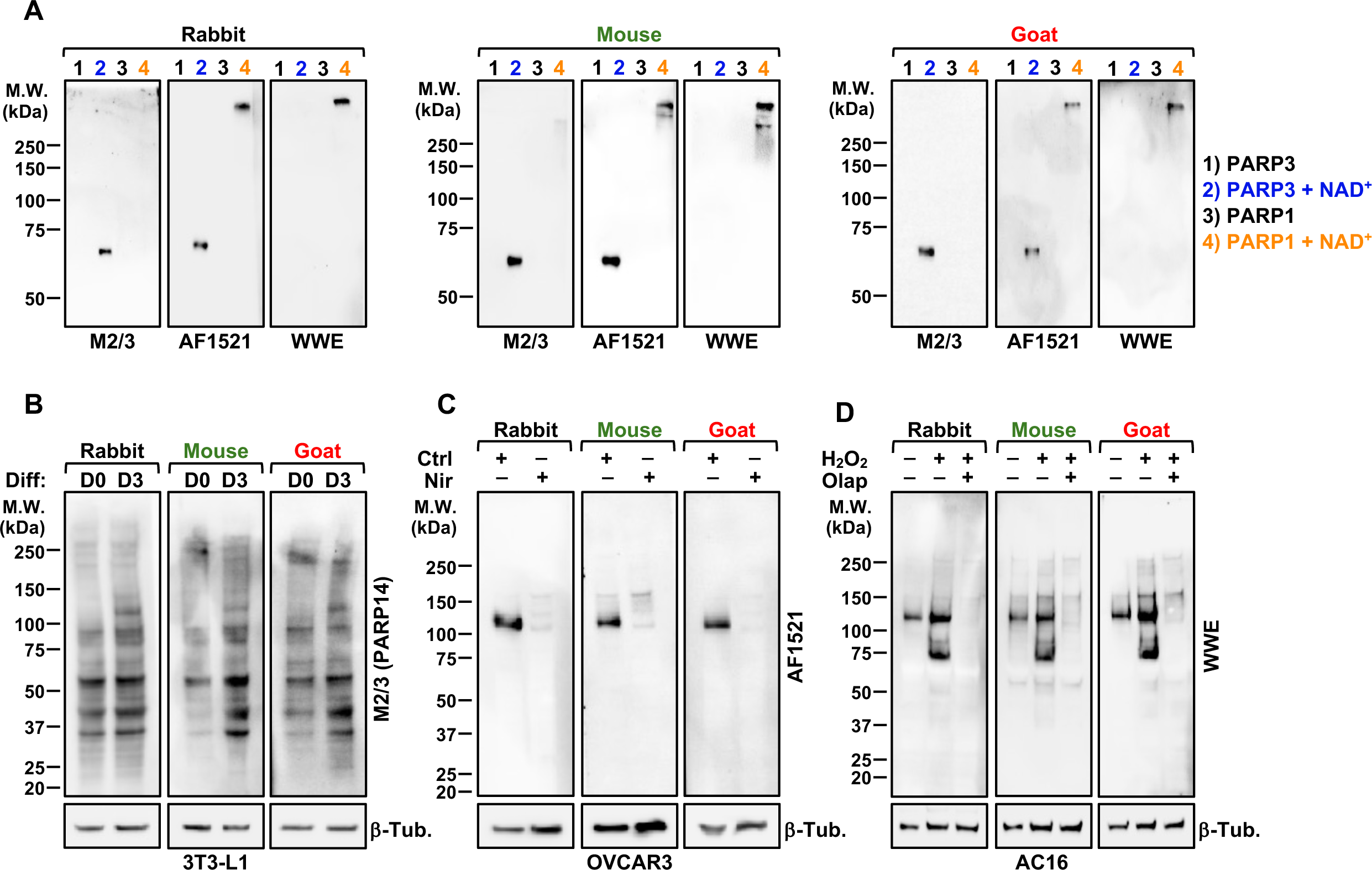
ADP-ribosylation specificity of new ARBD-detection reagents. **(A)** Immunoblot of mono- and poly (ADP-ribosyl)ated PARP proteins using ARBD-Fc fusion proteins for detection. Purified recombinant PARP1 or PARP3 were incubated with or without NAD^+^ to promote auto (ADP-ribosyl)ation. The yield was mono (ADP-ribosyl)ated PARP3 (*lane 2; blue*), or poly (ADP-ribosyl)ated PARP1 (*lane 4; orange*). The mono- and poly (ADP-ribosyl)ated PARP proteins were separated by SDS-PAGE, transferred to nitrocellulose membrane, and subjected to immunoblotting using the nine ARBD-Fc fusion proteins, as indicated. Each lane contained approximately the same number of terminal ADP-ribose units, to our best approximation. Molecular weight markers in kilodaltons (kDa) are indicated. **(B)** Immunoblot analyses of ADP-ribosylation in 3T3-L1 cells using ARBD-Fc fusion proteins. Cytosolic extracts were prepared from 3T3-L1 cells grown in culture and differentiated as described previously (Day 0/Undifferentiated;Day 3/Differentiated). Equal total protein levels were separated by SDS-PAGE, transferred to nitrocellulose membrane, and subjected to immunoblotting using the 3 Fc-species versions of M2/3P14-Fc. Molecular weight markers in kilodaltons (kDa) are indicated. **(C)** Immunoblot analyses of ADP-ribosylation in OVCAR3 cells using ARBD-Fc fusion proteins. Whole cell extracts were prepared from OVCAR3 cells grown in culture following treatment without or with 20 μM PARP inhibitor, nirparib. Equal total protein levels were separated by SDS-PAGE, transferred to nitrocellulose membrane, and subjected to immunoblotting using the 3 Fc-species versions of AF1521-Fc. Molecular weight markers in kilodaltons (kDa) are indicated. **(D)** Immunoblot analyses of ADP-ribosylation AC16 cells using ARBD-Fc fusion proteins. Whole cell extracts were prepared from AC16 cells grown in culture with out any treatment or following treatment with 5 mM H_2_O_2_ and 10 μM PARP inhibitor, olaparib. Equal total protein levels were separated by SDS-PAGE, transferred to nitrocellulose membrane, and subjected to immunoblotting using the 3 Fc-species versions of WWE-Fc. Molecular weight markers in kilodaltons (kDa) are indicated.

To explore the utility of the new reagents in biological systems, we performed immunoblotting in three different cell-based models. First, we analyzed mono ADP-ribosylation in 3T3-L1 cells grown in culture and differentiated as previously described *20*. Using the M2/3 (PARP14) detection reagent, we observe an increase in MARylation upon differentiation with all three species versions (Day 0 vs Day 3; Figure 3B; Table S3). Second, we analyzed poly ADP-ribosylation in OVCAR3 ovarian cancer cell line treated in culture with and without the PARP inhibitor (PARPi), niraparib. Using the AF1521 detection reagent, we observe a clear reduction in PARylation signal upon treatment with PARPi (Figure 3C). Third, we analyzed poly ADP-ribosylation in AC16 cardiomyoctes treated in culture with and without H_2_O_2_ and the PARPi, olaparib. Using the WWE detection reagent, we observe a clear increase in PARylation signal upon treatment with H_2_O_2,_ which can be abrogated by pretreatment with PARPi (Figure 3D). These results indicate that the detection reagents can recognize the expected specific forms of ADPR.

### Expansion of species-specific ARBD-Fc fusion proteins allows for multiplexing capabilities

Given the reagent limitations in the field of PARP biology, we set out to create tools that would allow for more multiplexed assessments that can help tease out complexities of ADPR biology in biological systems. Thus, we tested various co-stains and assayed by immunofluorescent immunocytochemistry and dual-color fluorescent Western blotting (Table S4). First, using OVCAR3 cell line treated with and without PARPi (veliparib or nirparib, as indicated), we co-stained to examine levels of PARylation and MARylation in cells by immunofluorescent immunocytochemistry. We observe a clear reduction in PARylation upon PARPi treatment, while MARylation remained largely unchanged (Figure 4A-B). We also used 3T3-L1 cells in a differentiation protocol. We observe PARylation decrease, while MARylation increases, in reponse to differentiation signals (Figure 4C).

**Figure 4.**
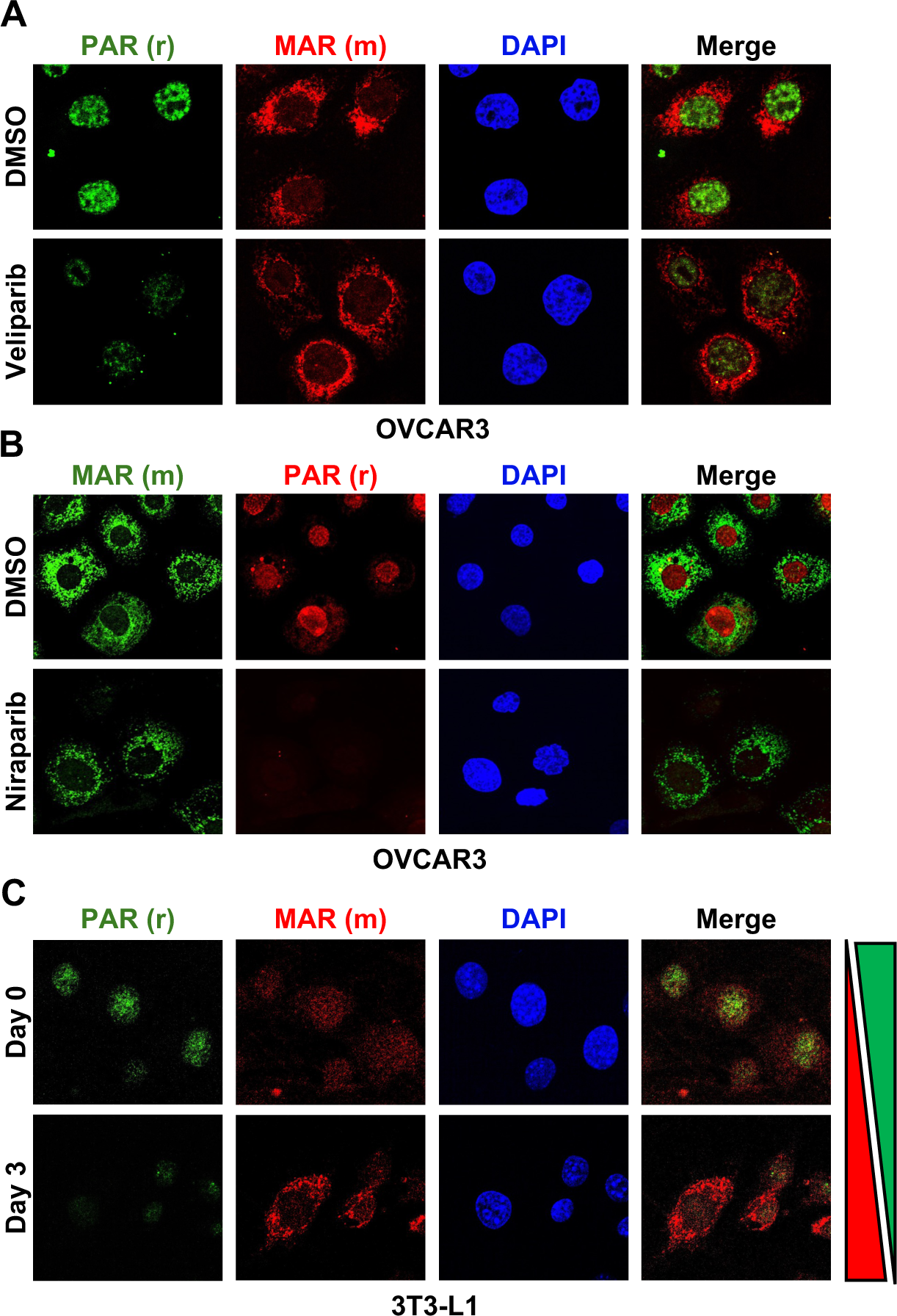
Testing new ARBD-detection reagents for fluorescent immunocytochemistry. **(A and B)** Immunofluorescent staining of ADP-ribosylation in OVCAR3 cells using ARBD-Fc fusion proteins. OVCAR3 cells grown in chamber slides were treated with DMSO or PARP inhibitors veliparib (A) or niraparib (B), as indicated. Following treatment, the cells were fixed with paraformaldehyde and co-immunostained for ADP-ribose using ARBD-Fc fusion proteins: Poly (ADP-ribosyl)ation (WWE-rabbit) and Mono (ADP-ribosyl)ation [M2/3 (PARP14)-mouse]. Different secondary fluorophores were used in each experiment. DAPI (blue) and merged images shown. Scale bar **(C)** Immunofluorescent staining of ADP-ribosylation in 3T3-L1 cells using ARBD-Fc fusion proteins. 3T3-L1 cells were grown in chamber slides and differentiated as described previously. Following the indicated time, the cells were fixed with paraformaldehyde and co-immunostained for ADP-ribose using ARBD-Fc fusion proteins: Poly (ADP-ribosyl)ation (WWE-rabbit) and Mono (ADP-ribosyl)ation [M2/3 (PARP14)-mouse]. DAPI (blue) and merged images shown. High PAR (green) level at Day 0 and high MAR (red) level at Day 3 is shown. Scale bar

Second, using the same 3T3-L1 differentiation protocol, we made whole cell extracts to examine various co-stains by dual-color fluorescent Western blotting. For example, we are able to successfully co-stain using the M2/3 (PARP14)-rabbit reagent with RPS6-mouse (Figure 5A; Table S4). Additionally, we demonstrate a clear overlap in signal using the goat and rabbit versions of WWE and AF152, respectively (Figure 5B). The signals are robust and comparable to what we usually observe using WWE-rabbit in an HRP-based detection system (Figure 5B; *far right panel*). All together, these results indicate that having the species variety in ARBD-Fc reagents is a valuable tool for studying ADPR biology.

**Figure 5.**
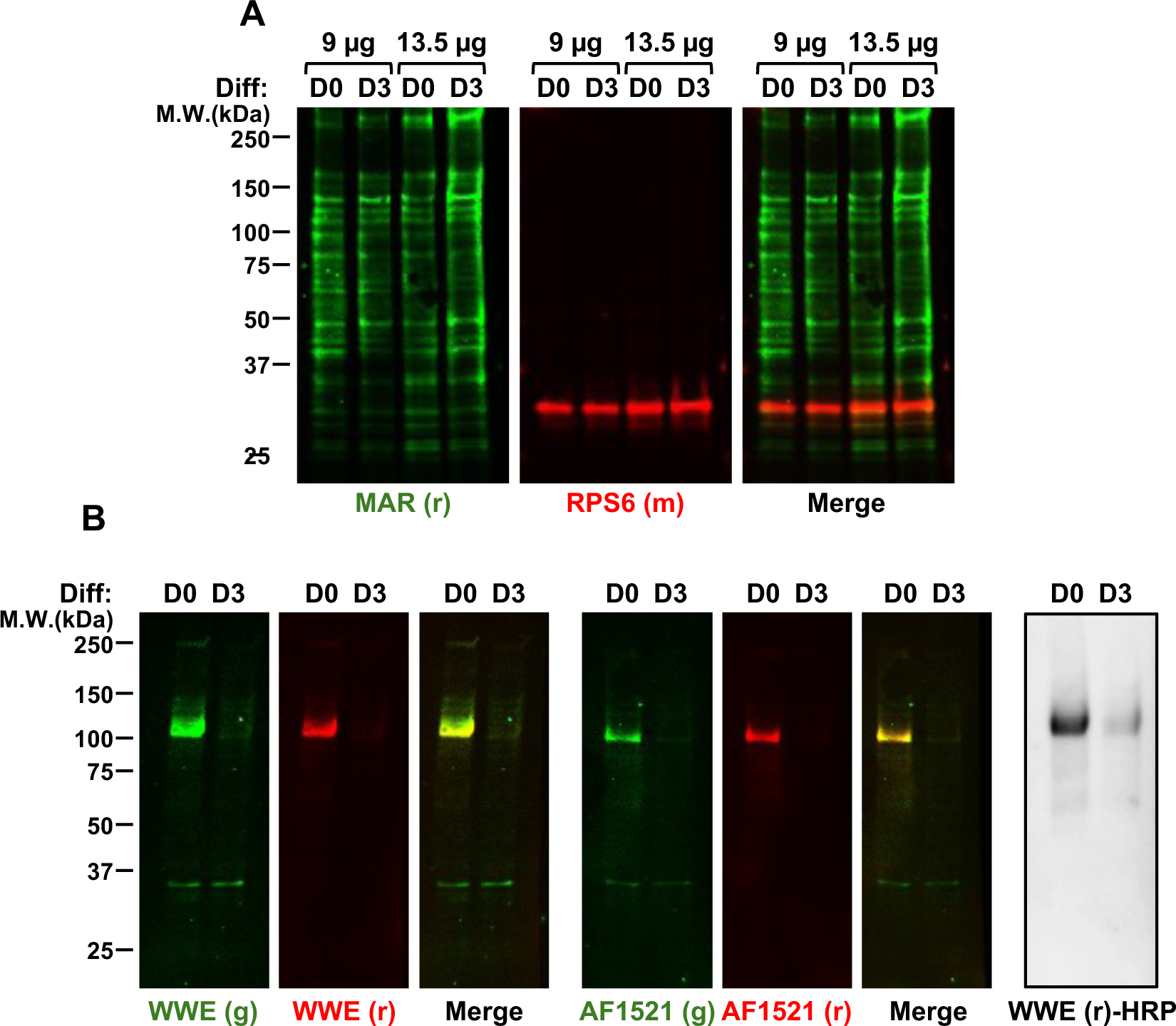
Testing new ARBD-detection reagents for dual-color fluorescent Western blotting. **(A)** Dual-color immunofluorescence analyses of ADP-ribosylation in 3T3-L1 cells using ARBD-Fc fusion proteins. Whole cell extracts were prepared from 3T3-L1 cells grown in culture and differentiated as described previously. Equal total protein levels were separated by SDS-PAGE, transferred to nitrocellulose membrane, and blotted for MAR: M2/3 (PARP14)-rabbit (green) and RPS6-mouse (red). **(B)** Dual-color immunofluorescence analyses showing overlap immunofluorescence signal (yellow) using WWE-goat (green) and WWE-rabbit (red) *(left panel*), AF1521-goat (green) and AF1521-rabbit (red) (*middle panel*), and immunoblotting using the WWE-rabbit and anti-rabbit HRP-conjugated antibody for comparison (*right panel*).

## Discussion

Herein, we have generated and characterized an expanded set of antibody-like ADP-ribose binding proteins in which these natural and specific ARBDs have been functionalized with the Fc region of mouse and goat immunoglobulin. In combination with the rabbit versions previously described *4*, we have created a more useful array of ADP-ribose detection reagents. Characterization of the new reagents indicates that they can be detected in a species-dependent manner and recognize specific ADP-ribose moieties. Importantly, these new immunological reagents can be used in various antibody-based assays by co-staining. The expansion of this tool will allow for more multiplexed assessments of the complexity of ADPRylation biology in many biological systems.

In order to study the complex and diverse forms and functions of ADPR, immunological tools used for its detection and enrichment are necessary, but still quite limited. Considerable efforts have been made to develop antibodies to detect ADP-ribose (MAR and PAR) *7-15*. The original rabbit versions of the antibody-like ADP-ribose binding reagents we previsouly generated have proven useful research tools in the field for the molecular recognition of various forms of ADP-ribose *4*. They are the first examples of functionalized ARBD-Fc fusion proteins that include all key features of a monoclonal antibody: 1) monospecificity, 2) binding to protein A and G, 3) binding to Ig-directed secondary antibodies, and 4) renewable production *4*. Most importantly, for the first time, these reagents allowed for the distinct recognition of mono- or oligo-ADPRylation, which had been largely overlooked by the widely-used anti-PAR monoclonal antibody 10H *7*. More recently, with improved methods for the chemical synthesis of ADPRylated susbtrates, coupled with phage display and SpyTag technology, the Matic laboratory has developed a new generation of site-specific, as well as broad-specificity antibodies to MAR *16, 17*.

The expansion of the ARBD-Fc toolkit by including fusions with mouse or goat Fc greatly enhance their utility in a wide range of immunological assays. Particularly, the new reagents allow for multiplexed assessment of MAR, OAR, and PAR, which is diverse, rich and complex, contributing to interesting biologies for ADPR in a variety of biological systems. The dual detection of MAR and PAR with these new reagents will also shed light on the interesting signaling interplay between these PTMs occurring simultaneously, in different subcellular compartments, in distinct biological processes, providing a level of precision that has yet to be attained in the field.

Hence, the continued development of tools and refinement of existing ones will undoubtedly enhance our understanding of the intricate biological functions of ADPRylation.

## Experimental Procedures

### Construction of plasmid vectors for bacterial expression of ADPR binding domain−Fc fusion proteins

The cDNA sequences encoding the Mouse and Goat IgG are listed in Supporting Information materials (Figure S1 and S2). The cDNA fragments were synthesized as gene blocks by Integrated DNA Technologies (IDT), and designed with appropriate restriction sites for cloning. The gene blocks were amplified using polymerase chain reaction (PCR). For the new constructs, the Rabbit-Fc region in the pET19b vector (Novagen) containing Macro2/3 (PARP14), Macro (AF1521), or WWE(RNF 146) *4* was excised using (1) the AgeI and EcoRI sites and replaced with PCR-amplified DNA encoding Mouse-Fc which was digested using AgeI and EcoRI or (2) the SalI and BamHI sites and replaced with PCR-amplified DNA encoding Goat-Fc which was digested using XhoI and BamHI. Sequences were verified using whole plasmid sequencing (Plasmidsaurus).

### Expression and purification of the antibody-like ADPR binding reagents in bacteria

#### Expression

The ADPR binding reagents were expressed in bacteria using the pET19b-based vectors described above. Because of expression issues, a slightly modified protocol was used described in Table S1. *E. coli* strain BL21(DE3) Rosetta2 pLysS was made competent using a CaCl_2_ protocol and transformed with the pET19b-based plasmids encoding one of the ADPR binding reagents described above. For 37°C induction, the transformed bacteria were grown in LB containing ampicillin and chloramphenicol at 37°C until the OD_595_ reached 0.4−0.6. Recombinant protein expression was induced by the addition of isopropyl β-D-1-thiogalactopyranoside (IPTG) for 4 h at 37°C. For 16°C induction, the transformed bacteria were grown in LB containing ampicillin and chloramphenicol at 37°C until the OD_595_ reached 0.2. The culture was then cooled to 16°C and grown to an OD_595_ of <0.8. Recombinant protein expression was then induced by IPTG for 18 h at 16°C. In all cases, the cells were collected by centrifugation, and the cell pellets were flash-frozen in liquid N_2_ and stored at −80°C.

#### Purification

The frozen pellets were thawed on wet ice and lysed by sonication in Ni-NTA lysis buffer [10 mM Tris-HCl (pH 7.5), 0.5 M NaCl, 0.1 mM EDTA, 0.1% NP-40, 10% glycerol, 10 mM imidazole, 1 mM phenylmethanesulfonyl fluoride (PMSF), and 1 mM β-mercaptoethanol]. The lysates were clarified by centrifugation at 20,000g using an SF14-50 rotor (Lynx6000, Sorvall) at 4°C for 30 min. The supernatant was incubated with 1 mL of Ni-NTA resin equilibrated in Ni-NTA equilibration buffer [10 mM Tris-HCl (pH 7.5), 0.5 M NaCl, 0.1% NP-40, 10% glycerol, 10 mM imidazole] at 4°C for 2-3 h with gentle agitation. The resin was collected by centrifugation at 4°C for 10 min at 800g, and the lysate (supernatant) was removed. The resin was washed four times with Ni-NTA wash buffer [10 mM Tris-HCl (pH 7.5), 1 M NaCl, 0.2% NP-40, 10% glycerol, 10 mM imidazole, and 1 mM PMSF]. The recombinant proteins were then eluted using Ni-NTA elution buffer [10 mM Tris-HCl (pH 7.5), 0.2 M NaCl, 0.1% NP-40, 10% glycerol, 500 mM imidazole, 1 mM PMSF, and 1 mM β-mercaptoethanol]. The eluates were collected by centrifugation (4°C for 5 min at 800 g) and dialyzed in Ni-NTA dialysis buffer [10 mM Tris-HCl (pH 7.5), 0.2 M NaCl, 10% glycerol, 10 mM imidazole, 0.1% NP-40, 1 mM PMSF, and 1 mM β-mercaptoethanol] (Table S2). The dialyzed proteins were collected and centrifuge at max speed (15,000 RPM) for 15 minutes in 4°C. Collected the clear supernatant and quantified using a Bradford protein assay (Bio-Rad), aliquoted, flash-frozen in liquid N_2_, and stored at −80 °C. To assess the purity and quality of each purified ADPR binding reagent, 1-2 μg of purified protein was subjected to sodium dodecyl sulfate−polyacrylamide gel electrophoresis (SDS−PAGE) and stained with Coomassie brilliant blue.

### In vitro auto(ADP-ribosyl)ation reactions with PARP1 and PARP3 to generate mono-, and poly(ADP-ribose) standards

Five hundred nanograms of purified recombinant mPARP1 or mPARP3 was incubated at 25°C in a 100 μL reaction volume [20 mM HEPES (pH 8.0), 5 mM MgCl_2_, 5 mM CaCl_2_, 0.01% NP-40, 25 mM KCl, 1 mM DTT, 0.1 mg/mL sheared salmon sperm DNA (Invitrogen, AM9680), and 0.1 mg/mL BSA (Sigma)] under the following conditions: (1) PARP1 with 250 μM NAD^+^ for 5 min for poly(ADP-ribose), and (2) PARP3 with 250 μM NAD^+^ for 30 min for mono(ADP-ribose). All reactions were stopped by the addition of one-third of a reaction volume of 4× SDS−PAGE loading buffer, followed by heating to 95°C for 5 min.

### Dot blotting and immunoblotting

#### Preparation of nitrocellulose blotting membranes for dot blotting and immunoblotting

For dot blotting, 500 ng purified proteins were spotted in 5 μL amounts to a nitrocellulose membrane and dried to bind the ADPR binding reagents to the membrane. For immunoblotting, the aliquots of purified proteins , PARP1 or PARP3 ADP-ribosylation reaction products, or 9 to 30 μg of nuclear or whole cell extract, were resolved on a 10% SDS-PAGE gel and transferred to a nitrocellulose membrane.

#### Dot blotting and immunoblotting

The membranes were blocked for 1 h at room temperature in Tris-buffered saline with 0.05% Tween (TBST) containing 5% nonfat dry milk. Primary antibodies and/or detection reagents (e.g., ADPR binding reagents; β-tubulin, Abcam ab6046) were diluted in TBST with 3% nonfat dry milk and incubated with membranes for 1 h at room temperature. As described in Table S3, after being extensively washed with TBST, the membranes were incubated with an appropriate HRP-conjugated secondary antibody (IgG Fc Goat anti-Rabbit, HRP, Invitrogen 31463; IgG Fc Rabbit anti-Mouse, HRP, Invitrogen 31455; IgG Fc Rabbit anti-Goat, HRP, Invitrogen 31433), and diluted in TBST with 3% nonfat dry milk for 1 h at room temperature. The membranes were washed extensively with TBST before chemiluminescent detection using SuperSignal West Pico substrate or femto substate (Thermo Scientific) and a ChemiDoc imaging system (Bio-Rad).

### Cell culture and treatments

3T3-L1 cells *21* cells were obtained from the American Type Cell Culture (ATCC, CL-173). They were maintained in DMEM (Cellgro, 10-017-CM) supplemented with 10% fetal bovine serum (Atlanta Biologicals, S11550) and 1% penicillin/streptomycin. For the induction of adipogenesis, the 3T3-L1 cells were grown to confluence and then cultured for 2 more days under contact inhibition. The cells were then treated for 2 days with an MDI adipogenic cocktail containing 0.25 mM IBMX, 1 μM dexamethasone, and 10 μg/ml insulin. Subsequently, the cells were cultured in medium containing 10 μg/ml insulin for the indicated times before collection *20*.

OVCAR3 cells were obtained from ATCC (HTB-161). The cells were cultured in RPMI 1640 (Gibco) supplemented with 10% fetal bovine serum (Sigma), 1% penicillin/streptomycin, and 1% glutamax. The cells were grown to ∼80% confluence, treated for 2 h with or without 20 μM PARP inhibitor, niraparib. Treated cells were gently washed and collected in ice-cold PBS and then pelleted by centrifugation at 450 g for 5 min.

AC16 human adult ventricular cardiomyocyte cells were obtained from ATCC (CRL-3568). The cells were cultured in DMEM/F-12, no phenol red (Gibco, 11039021) supplemented with 12.5% fetal bovine serum (Sigma, F8067), 1% penicillin/streptomycin, and 0.025 mg/ml Gentamicin (Gibco, 15710064). The cells can be cultured for at least 10 passages after initial thawing. AC16 cells were grown to ∼80% confluence, treated for 2 h with or without 10 μM PARP inhibitor, olaparib, and then treated for an additional 10 min with or without 5 mM H_2_O_2_.

For all cell lines, cell stocks were regularly replenished from the original stocks, verified for cell type identity, and confirmed as mycoplasma-free every three months using a commercial testing kit.

### Preparation of extracts from mammalian cells for immunoblotting

#### Preparation of whole cell extracts.

The cell pellets were resuspended in 1× lysis buffer [50 mM Tris (pH 7.5), 0.5 M NaCl, 1 mM EDTA, 1% NP40, 10% glycerol, 1 M DTT, 10 μM PJ34 (abcam, ab120981), 500 nM ADP-HPD (a PARG inhibitor; sigma) and 1× protease inhibitor cocktail (Roche)] and incubated for 30 min on ice with gentile vortexing and then centrifuge at maximum speed for 15 minutes at 4°C in microcentrifuge to remove the cell debris. The supernatant was collected as whole cell protein extracts. Protein concentrations for the whole cell extracts were determined using a Bradford protein assay (Bio-Rad). The extracts were aliquoted, flash-frozen in liquid N_2_, and stored at −80°C.

#### Preparation of nuclear extracts

The packed cell volume (PCV) was estimated, and the cell pellets were resuspended to homogeneity in 5× PCV of Lysis buffer [1x isotonic lysis buffer (10 mM Tris-HCl (pH 7.5), 2 mM MgCl_2_, 3 mM CaCl_2_, 0.3 M sucrose), 1 mM DTT, 1× protease inhibitor cocktail (Roche), 10 μM PJ34 (Sigma), and 500 nM ADP-HPD (a PARG inhibitor; sigma)] and incubated on ice for 15 min. NP-40 was added from a 20% solution in lysis buffer to a final concentration of 0.6%, and the mixture was vortexed vigorously for 10 s. The lysate was subjected to a short burst of centrifugation for 30 s at 11000 RPM in a microcentrifuge at 4°C to collect the nuclei. The pelleted nuclei were resuspended in 2/3x PCV of ice-cold nuclear extraction buffer C [20 mM HEPES (pH 7.6), 1.5 mM MgCl_2_, 0.42 M NaCl, 0.2 mM EDTA, 25% (v/v) glycerol, 1 mM DTT, 1× protease inhibitor cocktail, 10 μM PJ34 inhibitor, and 500 nM ADP-HPD] and incubated while being gently mixed for 30 min at 4°C. The mixture was subjected to centrifugation at maximum speed in a microcentrifuge for 10 min at 4°C twice to remove the insoluble material. The supernatant was collected as the soluble nuclear extract. Add equal volume of buffer B [20 mM HEPES (pH 7.6), 1.5 mM MgCl_2_, 0.2 mM EDTA, 1 mM DTT, 1× protease inhibitor cocktail, 10 μM PJ34 inhibitor, and 500 nM ADP-HPD]. NP-40 was added from a 20% solution in lysis buffer to a final concentration of 0.5%. Protein concentrations for the nuclear extracts were determined using a Bradford protein assay (Bio-Rad). The extracts were aliquoted, flash-frozen in liquid N_2_, and stored at −80°C.

Aliquots of nuclear or whole cell extracts were mixed with one-third of a volume of 4× SDS loading buffer [200 mM Tris-HCl (pH 6.8), 8% SDS, 40% glycerol, 4% β-mercaptoethanol, 50 mM EDTA, and 0.08% bromophenol blue], followed by heating to 70°C for 10 min. The cell extracts were subjected to immunoblotting as described above.

### Immunofluorescent immunocytochemistry

OVCAR3 cells were grown on chamber slide (Invitrogen), treated for 2 h with or without 20 μM PARP inhibitor (veliparib, niraparib). 3T3-L1 cells were seeded into 4-well chambered slides (Thermo Fisher, 154534) and were grown and differentiated as described previously *20*. The cells were washed twice with PBS on ice and fixed with 4% paraformaldehyde (EMS) at room temperature for 15 min. The chamber slides were washed with ice-cold PBS and permeabilized with permeabilization buffer (1x PBS, 0.5% Triton X-100) for 5 min. The fixed cells on the chamber slides were washed with PBS and blocked with blocking solution for 1 h at room temperature. After being blocked, the samples were incubated with 20 μg/mL ADPR binding reagent in blocking solution overnight at 4°C. The samples were washed with PBST and incubated with fluorophore-conjugated secondary antibodies (Goat anti-Rabbit IgG (H+L) Alexa Fluor 594, A-11012; Goat anti-Rabbit IgG (H+L), Alexa Fluor 488, A-11008; Goat anti-Mouse IgG (H+L), Alexa Fluor 594, A-11005; Goat anti-Mouse IgG (H+L), Alexa Fluor 488, A-11001) diluted 1:500 in PBST for 1 h at room temperature in the dark. The samples were then washed with PBST, the coverslips were placed on cells coated with VectaShield Antifade Mounting medium with DAPI (Vector Laboratories), and image was visualized using Nikon confocal microscope.

### Dual-color fluorescent Western blotting

The membranes were blocked for 1 h at room temperature in Tris-buffered saline with 0.05% Tween (TBST) containing 3% nonfat dry milk (Table S4). Primary antibodies and/or detection reagents (e.g., ADPR binding reagents; RPS6, Santa Cruz sc-74459) were diluted in TBST with 3% nonfat dry milk and incubated with membranes for 1 h at room temperature. After being extensively washed with TBST, the membranes were incubated with secondary antibodies (IRDye 800CW Donkey anti-Rabbit IgG Secondary Antibody, Li-Cor 926-32213; IRDye 680RD Donkey anti-Rabbit IgG Secondary Antibody, Li-Cor 926-68073; IRDye 680RD Donkey anti-Mouse IgG Secondary Antibody, Li-Cor 926-68072; IRDye 800CW Donkey anti-Goat IgG Secondary Antibody, Li-Cor 926-32214) diluted in TBST with 3% nonfat dry milk for 1 h at room temperature. The membranes were washed extensively with TBST. The image was developed using ChemiDoc imaging system (Bio-Rad).

## Data Availability

All data and reagents presented within this article are available upon request. The generated plasmids will be available through EMD Millipore.

## Supporting Information

Supporting information includes additional figures and Table S1-S4 and can be found with this article online. [See the Supporting Information file.]

## Acknowledgements

We thank members of the Kraus lab Aarin Jones and MiKayla S. Stokes for help in cloning new constructs; MiKayla S. Stokes, Palak Ahuja, Marwa W. Aljardali, and Xinrui Tan for providing cell lysates for screening the detection reagents; Dan Huang for providing purified protein preparations and Charles W. Renshaw for help with purifications and immunoblotting assays.

## Author Contributions

W.L.K. - conceptualization, funding acquisition; S.C., C.V.C. - data curation, formal analysis, methodology, validation, writing-original draft. W.L.K., C.V.C. - supervision, writing-review and editing, project administration.

## Funding and Additional Information

This work was supported by grants from the NIH/National Cancer Institute (R01 CA251943 to W.L.K.), NIH/National Institute of Diabetes and Digestive and Kidney Diseases (R01 DK069710 to W.L.K.), Cancer Prevention and Research Institute of Texas (CPRIT; RP220325 to W.L.K.), U.S. Deparment of Defense Ovarian Cancer Research Program (DOD-OCRP; OC200311 to W.L.K.), and funds from the Cecil H. and Ida Green Center for Reproductive Biology Sciences Endowment to W.L.K. The content is solely the responsibility of the authors and does not necessarily represent the official views of the National Institutes of Health.

