## Supplemental Materials for "Development and Characterization of Recombinant ADP-Ribose Binding Reagents that Allow Simultaneous Detection of Mono and Poly ADP-Ribose"

**Chiu *et al.* (2024)**

This document contains the following supporting information: Page

### SUPPORTING FIGURES

#### Figure S1. Cloning of mouse (*Mus musculus*) Fc

**Original Fc amino acid sequence of laboratory mouse (*Mus musculus*)**
GCKPCICTVPEVSSVFIFPPKPKDVLTITLTPKVTCVVVDISKDDPEVQFSWFVDDVEVHT
AQTQPREEQFNSTFRSVSELPIMHQDWLNGKEFKCRVNSAAFPAPIEKTISKTKGRPKAP
QVYTIPPPKEQMAKDKVSLTCMITDFFPEDITVEWQWNGQPAENYKNTQPIMDTDGSYF
VYSKLNVQKSNWEAGNTFTCSVLHEGLHNHHTEKSLSHSPGK

*GenBank AAK53870.1:* This amino acid sequence was used to generate cDNA sequence for cloning.

**AgeI-Mouse Fc-EcoRI-BamHI gene block sequence (IDT)**
AGGTCTTTGAGAGGAGTCTTGGGTCG**ACCGGT**AGTTCAATGGGATGCAAGCCATGCATCTGCAC
CGTTCCCGAAGTGTCATCGGTGTTCATTTTCCCTCCCAAGCCGAAGGATGTTCTTACTATCACC
CTTACGCCTAAAGTTACCTGTGTTGTCGTCGATATTTCAAAAGACGACCCAGAGGTGCAGTTTT
CGTGGTTTGTAGATGATGTTGAAGTTCATACCGCCCAGACGCAACCGCGTGAAGAACAGTTTAA
TTCCACCTTTCGCTCCGTATCTGAGCTTCCTATCATGCACCAGGACTGGCTTAATGGTAAGGAA
TTTAAATGCCGCGTGAACAGTGCTGCATTCCCCGCTCCAATTGAAAAGACAATCAGTAAGACTA
AAGGGCGTCCGAAAGCCCCGCAGGTTTACACAATTCCTCCACCGAAAGAACAAATGGCGAAGGA
TAAAGTATCCCTGACTTGTATGATTACAGACTTTTTCCCCGAAGACATTACTGTAGAGTGGCAG
TGGAATGGACAGCCCGCGGAGAATTACAAAAATACCCAACCTATCATGGATACAGATGGTAGCT
ACTTCGTGTACTCTAAGTTGAATGTTCAGAAGTCCAACTGGGAAGCCGGAAATACTTTTACGTG
CTCTGTTTTACACGAGGGCCTTCATAACCATCATACCGAGAAGTCATTGTCGCATTCCCCAGGT
AAGTAATAATA**GAATTCGGATCC**GGCTGCCAATTGTAACAAA

Primers used for PCR amplification of gene fragment:
AgeI-Mouse-Forward 5'-TTGAGAGGAGTCTTGGGTCGACCGG-3'
EcoRI-Mouse-Reverse 5'-TACAATTGGCAGCCGGATCCGAATT-3'

**Figure S2. Cloning of goat (*Capra hirus*) Fc**

**Original Fc amino acid sequence of domestic goat (*Capra hirus*)**
EPCQCPKCPEPLGGLSVFIFPPKPKDTLTISGTPEVTCVVVDVGQDDPEVQFSWFMDNVE
VHTARTTPREEQFNSTFRVVSALPIQHKDWLQGKEFKCKVHNEGLPAPIIRTISRAKGQA
REPQVYVLAPPREELSKSTLSVTCLITGFYPEEVDVEWQRDGQPESEDKYHTAPPQLDA
DGSYFLYSRLRVNKSSWQEGDTYTCAVMHEALRNHYKEKSISKSPGK

*GenBank KAJ1069780.1:* This amino acid sequence was used to generate a codon optimized cDNA sequence for cloning using the (hinge+CH2+CH3) region.

Ref: Schwartz JC, Philp RL, Bickhart DM, Smith TPL, Hammond JA (2018) The antibody loci of the domestic goat (Capra hircus). *Immunogenetics* 70:317-326.

**XhoI-Goat Fc-EcoRI-BamHI codon optimized gene block sequence (IDT)**
TAAGCACTAGACGG**CTCGAG**CGGTAGTTCACCTTGTCAATGTCCAAAATGCCCCGAACCGCTGG
GCGGTCTTTCCGTGTTCATCTTTCCGCCTAAGCCGAAGGACACTTTGACCATTAGTGGCACCCC
GGAGGTTACGTGCGTTGTGGTGGACGTTGGTCAAGATGATCCGGAGGTCCAGTTCAGCTGGTTC
ATGGATAATGTTGAGGTGCACACTGCCCGCACCACCCCGCGTGAAGAGCAGTTCAACTCCACCT
TTCGTGTTGTCAGCGCATTACCGATTCAGCATAAAGACTGGCTGCAGGGTAAAGAATTTAAGTG
CAAAGTTCACAACGAAGGCCTGCCGGCACCGATCATCCGCACCATCTCTCGTGCGAAGGGTCAA
GCTCGTGAACCGCAGGTTTATGTTCTGGCGCCACCGCGTGAGGAACTGTCGAAGTCTACGCTGA
GCGTAACGTGCCTGATTACCGGTTTTTATCCGGAGGAAGTCGACGTGGAATGGCAGCGTGACGG
CCAACCGGAGAGCGAAGATAAATACCATACCGCGCCACCGCAACTGGACGCGGATGGTAGCTAC
TTCTTGTACAGCCGTCTGCGCGTGAATAAAAGCAGCTGGCAAGAGGGCGATACCTATACCTGTG
CTGTGATGCATGAAGCGTTGCGCAACCACTACAAAGAGAAAAGCATTTCCAAGTCCCCGGGTAA
GTGATAA**GAATTCGGATCC**GGCTGCTAAGCA

Primers used for PCR amplification of gene fragment:
XhoI-Goat-Forward 5'-CTAGACGGCTCGAGCGGTAGTTCAC-3'
BamHI-Goat-Reverse 5'-GCTTAGCAGCCGGATCCGAATTCT-3'

**Table S1. Summary of Conditions for Protein Expression in Bacteria**

| Domain | Fc-Species | IPTG induction 37°C | IPTG induction 16°C | Incubate with Ni-NTA |
|---|---|---|---|---|
| M2/3 (PARP14) | Rabbit | | *0.1 mM O/N | 2 hr |
| | Mouse | | **0.5 mM O/N | 2 hr |
| | Goat | | 0.1 mM O/N | 2 hr |
| Macro (AF1521) | Rabbit | 0.5 mM, 4hr | | 3 hr |
| | Mouse | | **0.5 mM O/N | 3 hr |
| | Goat | | 0.1 mM O/N | 3 hr |
| WWE (RNF146) | Rabbit | 0.5 mM, 4hr | 0.5 mM O/N | 2.5-3 hr |
| | Mouse | | 0.1 mM O/N | 2.5-3 hr |
| | Goat | | 0.1 mM O/N | 2.5-3 hr |
| Alternate [IPTG]: *0.2 mM** 0.1 mM | | | | |

**Table S2. Summary of Dialysis Conditions for Protein Purification**

| Domain | Fc-Species | Tubing membrane | Dilution |
|---|---|---|---|
| M2/3 (PARP14) | Rabbit | a,b | Dilute 2x-3x |
| | Mouse | a,b | |
| | Goat | a | |
| Macro (AF1521) | Rabbit | b | |
| | Mouse | a,b | Dilute 2x-3x |
| | Goat | a | |
| WWE (RNF146) | Rabbit | a,b | Dilute 2x-3x |
| | Mouse | a,b | Dilute 2x-3x |
| | Goat | a,b | Dilute 2x-3x |
| Dialysis tubing: a) 10K MWCO, 16 mm, SnakeSkin™ Dialysis Tubing<br>b) 45 mm MCO 3,500 Tubing | | | |

**Table S3. Summary of Conditions for Western Blotting**

| Domain | Fc-Species | Blocking Solution | µg of reagent in 5 ml blocking solution | HRP reagent |
|---|---|---|---|---|
| M2/3 (PARP14) | Rabbit | 5% BSA in TBST | 10 µg | Pico |
| | Mouse | 5% BSA in TBST | 10 µg | Pico |
| | Goat | 5% BSA in TBST | 20 µg | Femto |
| Macro (AF1521) | Rabbit | 3% milk in TBST | 10 µg | Pico |
| | Mouse | 3% milk in TBST | 20 µg | Pico |
| | Goat | 3% milk in TBST | 20 µg | Femto |
| WWE (RNF146) | Rabbit | 3% milk in TBST | 10 µg | Pico |
| | Mouse | 3% milk in TBST | 10 µg | Pico |
| | Goat | 3% milk in TBST | 20 µg | Femto |

**Table S4. Summary of Conditions for Dual-Color Fluorescent Western Blotting**

| Co-Stain | Blocking Solution | Amount in 5 mL blocking solution | LiCor |
|---|---|---|---|
| M2/3 (PARP14)- Rabbit<br>RPS6- Mouse | 3% BSA in TBST | 10 µg<br>1:3000<br>5% BSA in TBST | 1:25000<br>3% BSA in PBST |
| WWE (RNF146)- Rabbit<br>WWE (RNF146)- Goat | 3% milk in TBST | 10 µg<br>20 µg | 1:25000<br>3% milk in TBST |
| Macro (AF1521)- Rabbit<br>Macro (AF1521)- Goat | 3% milk in TBST | 10 µg<br>20 µg | 1:25000<br>3% milk in TBST |
